# A small H_2_O-soluble ingredient of royal jelly lower cholesterol levels in liver cells by suppressing squalene epoxidase

**DOI:** 10.1101/2022.06.14.496166

**Authors:** Chi Wang, Zhen-yu Jiang, Jing Wang, Jia-xin Liu, Yuan-yuan Nian, Lixia-Liu, Tong Dang, Xian-mei Meng

## Abstract

Excessive cholesterol in the liver is harmful for our health and may cause many diseases, such as fatty liver disease. Many studies in human and animal models have reported that royal jelly (RJ) can be used to treat atherosclerosis. However, the real mechanisms behind this action is unclear. In this study, we investigated the effectivity of RJ on gene expression of squalene epoxidase (SE) a major enzyme involved in cholesterol biosynthesis in HepG2 cells. We found that the expression of SE was decreased in response to RJ treatment. We also found that the origin of the RJ affected its strength. To find out the active ingredient of RJ in cholesterol suppression, we separated RJ into two parts based on the molecular weights using ultrafiltration membrane. We found that the fraction < 10kDa from RJ had comparable effect on SE expression, especially its water-soluble part. Taken together, we think RJ suppresses cholesterol by decreasing SE gene expression in liver. The active ingredient of RJ in this action is < 10kDa in water-soluble form.

## Introduction

Fatty liver disease is a clinic-pathologic syndrome caused by excess fat storage and steatosis of liver cells. It can occur in different races, ages, and sexes, but in China it is most common in the population at age from 40 to 49 ^[1,2]^. The prevalence rate among Chinese adults is currently as high as 15% to 20%, and there is a trend of morbidity to increase and for age to go down ^[3-7]^. In clinical, fatty liver disease is usually classified into non-alcoholic fatty liver disease (NAFLD) and alcoholic fatty liver disease (ALD) based on the history of alcohol consumption. Regardless, the consequences can be severe.

Three quarters of the ALD patients have an hepatomegaly and 30% of them has developed liver fibrosis and liver cirrhosis, or even liver cancer at some point ^[8]^. Some patients also have gastrointestinal symptoms, such as persistent right upper quadrant pain, significant loss of appetite, and severe diarrhea, leading to acute liver failure ^[9]^. NFALD can be as bad as ALD. After the intermediate-stage of the disease, some symptom such as liver pain, bloating, fatigue, poor appetite, and other symptoms of chronic hepatitis may occur. These symptoms also indicate an intermediate stage in the progression to cirrhosis and hepatocellular carcinoma. The NFALD deaths are usually resulted due to cardiovascular disease, liver cancer, and liver failure, all of which occur in the middle stages of Steatosis ^[10]^. Since the disease has almost no symptoms in its early stages, patients often die without any warning. In China, fatty liver disease has gradually replaced Hepatitis B becoming the first place of liver diseases. This is not sub-health but a disease. Although the real mechanisms of fatty liver disease remain unclear, increasing evidence shows a correlation between excessive cholesterol and fatty liver disease. Indeed, patients with fatty liver disease do have much higher cholesterol levels than the general population. Animal models also support this notion ^[11]^. Logically, lowering cholesterol levels in the liver should be a great way to prevent or cure fatty liver disease.

Cholesterol is a derivative of cyclopentane polyhydroxy phenylene and is synthesized mainly in the liver. Only 20% of cholesterol is obtained from food or other sources. The right amount of cholesterol is harmless to our body. It is essential component of the cell membranes, bile acids, vitamin D and steroids. However, elevated cholesterol levels can lead to a variety of diseases, such as coronary heart disease. The higher the total serum cholesterol, the greater the risk of arteriosclerosis. For every 1% reduction in total serum cholesterol, the risk of coronary heart disease can be reduced by 2% ^[12]^. There are many laboratory studies done to show that excessive cholesterol contributes to the development of fatty liver disease ^[11,13]^. Some studies have shown that the accumulation of cholesterol can directly damage the integrity of Mitochondria and endoplasmic reticulum membrane, trigger mitochondrial oxidative damage and stress, promote the formation of toxic oxidative sterols, and indirectly induce fat metabolism dysfunction ^[14]^. High levels of cholesterol have an indirect effect on liver health through Low-density Lipoprotein (LDL). Because the liver itself can only synthesize fat but not store it, the synthetic triglyceride is combined with cholesterol, phospholipids, etc. to very low-density Lipoprotein (VLDL) into blood and transported to extrahepatic tissues for storage or utilization. VLDL, however, can easily be metabolized and become LDL by excessive cholesterol. It is well known that LDL not only causes arteriosclerosis plaque to form, causing blood vessels to become inelastic and narrow; it also causes impaired secretion and synthesis of VLDL, which prevents fat from being excreted in time after it synthesized in the liver, this causes a lot of fat to accumulate in the liver, which then forms fatty liver. Even the accumulation of LDL may contribute to liver inflammation and fibrosis ^[14,15]^.

Royal Jelly (RJ) is the secretion of the pharyngeal gland of young worker bees that to fed and nurture larvae for the development. It is enriched with proteins and small peptides, also containing fats, sugars, acetylcholine-like substances, and hormones that human needs. It has been shown that the average worker bees fed by RJ is about twice as large as the average worker bees ^[16]^. RJ has high nutritional and medicinal values, such as lowering blood lipids, lowering cholesterol, preventing arteriosclerosis, etc ^[17-20]^. The function of RJ for lowering cholesterol and systolic blood pressure were reported 30 years ago ^[21]^. Some studies have shown in mice that the RJ up-regulated 148 genes and down-regulated 119 genes by > 1.8 fold compared to control. In another word 2.7% of the 10,000 genes were changed in expression in response to RJ. In liver genes, RJ increased the expression of fatty acid transporter 3(Fatp3), but decreased sterol regulatory element binding protein (SREBP)-1, and increased the expression of LDL receptor, but reduced squalene epoxidase (SE) ^[23]^. LDL receptors are involved in cholesterol uptake and transport, and SE is an essential enzyme directly involved in cholesterol biosynthesis in the liver. Without SE, cholesterol can not be produced in the liver.

However, animal experiment alone is inadequate to claim that RJ suppresses cholesterol production. After consumption by the animal, RJ is broken down into many small molecules, and which one of them is responsible for the experimental result is still a question. This study is to fill the gap using HepG2 cell line.

## Materials and Methods

### Cell culture

Human hepatoma HepG2 cell lines were purchased from the WHELAB cell company (Shanghai, China). Cells were cultured in growth medium which including low glucose-Dulbecco’s modified Eagle’s medium (LG-DMEM, Lining, Shanghai, China) supplemented with 10% heat-inactivated fetal bovine serum (FBS, Yuanye bio, Shanghai, China) and 1% 100 U/mL penicillin/streptomycin (Gibco, Scotland, UK). All the cells were incubated at 37°C under an atmosphere containing 5% CO_2._ HepG2 cells were detached by treatment at 37°C with 0.25 % trypsin– EDTA solution (Yuanye bio, Shanghai, China).

### RJ treatment

When the number of cells got confluence (about 80% of culture dish) during cell culture, the cells were detached with 0.25% trypsin-EDTA solution and diluted with growth medium to cell suspension of 2.5 × 10^5^ cells. Remove cell suspension 2 mL to per 35mm culture dish for 3 hours incubation, let cells fully attach to the bottom of the dish. The RJ treatment will be started after three hours incubation. The HepG2 cell line was exposed to various RJ added growth medium for 16 hours. The concentration of RJ added growth medium were (2.5mg/mL’5mg/mL’10mg/mL and 20 mg/mL). The control group was treated by growth medium.

### RNA extraction and Reverse transcription

After 16 hours treatment, RNA extraction experiments will be carried out by using the RNeasy Plus Mini Kit (QIAGEN, Germany). Using 0.25% trypsin-EDTA solution 0.75 mL detached cells from each 35 mm culture dish and washed with growth medium. Cell suspension was collected in a 1.5 ml centrifuge tube. The collected cell suspension was centrifuged at the rate of 3000rmp in a 4 °C for 5 minutes. After that, the supernatant was removed, the cells were dispersed by repeatedly tapping the bottom of centrifuge tube with the finger. 350μL Buffer RLT Plus reagent (one of reagent of the Mini kit) was added into tube, shaken and mixed. The cells mixed with the reagent were moved into the Qiashredder Tube (the tube in the Kit) and centrifuged at the rate of 15000RMP for 2 minutes for crushing the cells. After centrifugation, added equal volume of 70% alcohol and mixed sufficiently. Then the mixture was centrifuged at 8000g for 15 seconds. After emptying the filtrate, 700 μL Buffer RW1 reagent (one reagent of Mini kit) was added and centrifuged at a rate of 8000g for 15 seconds. After emptying the filtrate, 500 μ L Buffer RPE reagent (one reagent of Mini kits) was added and centrifuged at a rate of 8000g for 15 seconds to empty the filtrate and repeat the step twice. Dry up the RNA by centrifuging at a rate of 15,000 RMP for 1 minute. Then 100 μ L RNase-free water was added into the tube and centrifuging at a rate of 8000g for 1 minute. Then 100 μL RNA solution was obtained.

After the RNA solution obtained, the reverse transcription experiment was carried out. The concentration of RNA solution was measured by double beam UV photometer (PERSEE TU-1900, China) with 30 times TE buffer dilution (Baili Bio, Shanghai, China). According to measured concentration, obtained RNA solution was diluted to 200ng/10μL as Table 1. The RNA mixture was put into the Type TC gene amplifying apparatus(Bori TC-96/G/A(b)B)for cycle reaction (37°C, 15 minutes and then 85 °C for 5 seconds). Continuously added Master mix solution (Table 2) 10 μL for cycle reaction. After reaction, 20μL complementary DNA (cDNA) solution was obtained.

**Table 1.**
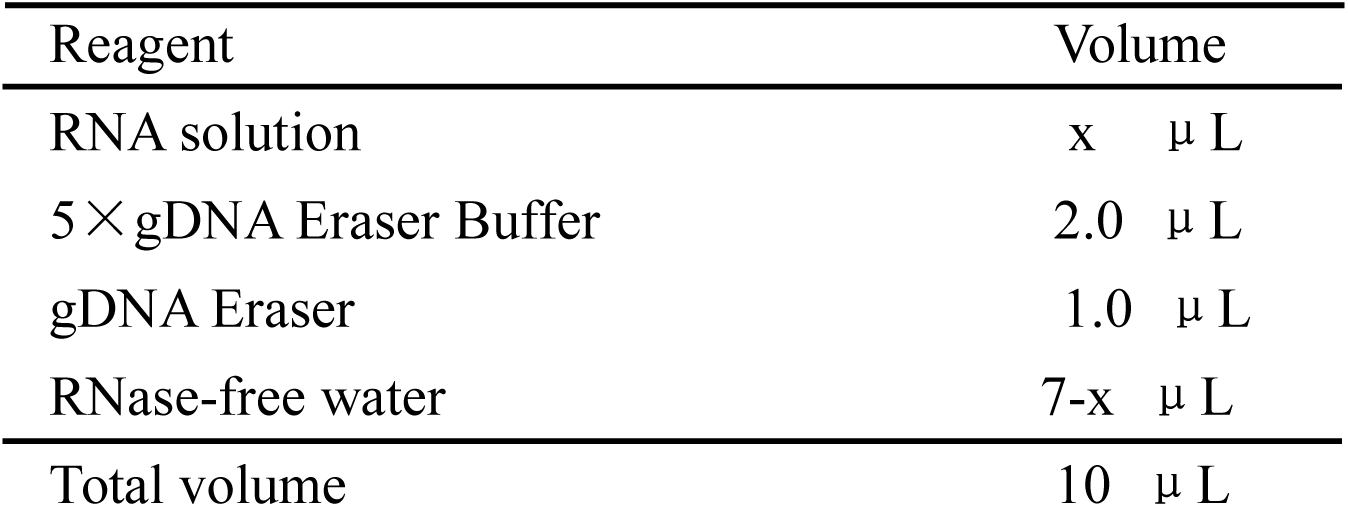

| Reagent | Volume |
| --- | --- |
| RNA solution | x $\mu$ L |
| 5×gDNA Eraser Buffer | 2.0 $\mu$ L |
| gDNA Eraser | 1.0 $\mu$ L |
| RNase-free water | 7-x $\mu$ L |
| Total volume | 10 $\mu$ L |

**Table 2.**

| Reagent | Volume |
| --- | --- |
| 5×PrimeScript Buffer 2 (for Real Time) | 4 $\mu$ L |
| PrimeScript RT Enzyme Mix I | 1 $\mu$ L |
| RT Primer Mix | 1 $\mu$ L |
| RNase Free water | 4 $\mu$ L |
| RNA solution | 10 $\mu$ L |
| Total volume | 20 $\mu$ L |

### Real-Time Polymerase Chain Reaction

Total cDNA was absorbed from RJ treated HepG2 cells by using the RNeasy Plus Mini Kit (QIAGEN, Germany). The reverse transcription reaction was performed using ExScript RT Reagent Kit (Takara, Tokyo, Japan). The primers used in this study were as follows:

glyceraldehyde-3-phosphate dehydrogenase (GAPDH):

Forward primer, 5′-AAATGGTGAAGGTCGGTGTG-3′;

Reverse primer, 5′-TGAAGGGGTCGTT GATGG-3′.

Squalene epoxidase (SE):

Forward primer, 5′-TGTCAGAAACCAACCAAGTGCAG-3′;

Reverse primer, 5′-TCCTTGTATTGCACGCCGATTA-3′.

All assays were performed using an ABI Prism 7500 Sequence Detection System (Applied Biosystems, Foster City, CA). Reactions were performed in a reaction mixture consisting of SYBR Premix Ex Taq (Takara, Tokyo, Japan) and primers with cDNA in a volume of 25 mL (Table3). The RT-PCR conditions were as follows: an initial step of 30 sec at 95°C, followed by 40 cycles of 5 sec at 95 °C and 30 sec at 60°C. The average expression level relative to the control group was determined after normalization with respect to the GAPDH in each sample.

**Table 3.**

| Reagent | Volume |
| --- | --- |
| TB Green Premix Ex Taq (Tli RNaseH Plus) (2×) | 12.5 $\mu$ L |
| PCR Forward Primer (50 $\mu$ M) | 0.5 $\mu$ L |
| PCR Reverse Primer (50 $\mu$ M) | 0.5 $\mu$ L |
| RNase-free water | 9.5 $\mu$ L |
| cDNA (25 times diluted by RNase-free water) | 2.0 $\mu$ L |
| Total volume | 25 $\mu$ L |

### Statistic analysis

The data of dual-lucifierase reporter and RT-PCR processed by T.TEST of Excel 2019. The results are presented as means –standard deviation. Furthermore, p less than 0.05 was regarded as statistically significant, p less than 0.01 was highly significant, and p larger than 0.05 was no significant.

## Results

### Gene expression of SE decreasing by RJ

To investigate whether RJ can decrease the gene expression of SE in liver, we incubated Human hepatoma cells (HepG2) with the RJ solution at the concentration of 2.5 mg/mL, 5.0 mg/mL, 10.0 mg/mL, for 16 hours, respectively. Total RNA of the cells was then extracted for the reverse transcription to produce complementary DNA (cDNA), which was used as the templates to determine SE expression by RT-PCR. No significant difference was found between group 2.5 mg/mL and the control, but in the rest of the treatment groups SE expression was decreased significantly (Fig 1a) indicating that RJ suppresses SE expression. When the concentration of royal jelly exceeds 20 mg/mL, a great number of cells died indicating that 2.5 ∼ 20 mg/mL is optimal. There was no significant difference between 10 and 20 mg/mL suggesting a possible threshold of decreasing cholesterol by RJ.

**Fig. 1.**
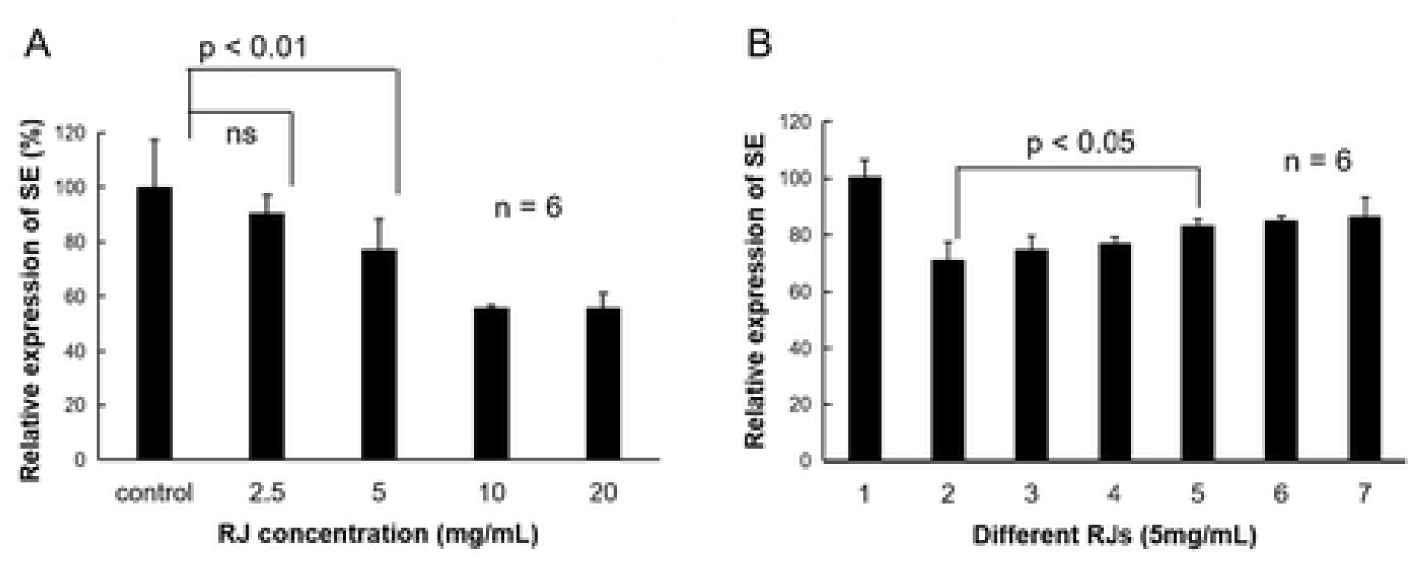
RJ directly decreasing SE gene expression in HepG2 cells, and different region collected RJ showed different effectiveness of suppression. (A) Representative Real-time polymerase chain reaction(RT-PCR) showing the expression changes of SE in the presence of 5 (RJ concentration: 5 mg/mL) and 10 (RJ concentration: 10 mg/mL). Vertical axis represent as the value of SE expression divided by the value of GAPDH expression. Horizontal axis represent as concentration of RJ. Control group was growth medium only. RT-PCR analysis shows that 5 (5 mg/mL) and 10 (10 mg/mL) significantly decrease mRNA expression of squalene epoxidase (SE) in HepG2 cells (p < 0.01). (B) The results of RT-PCR showing the expression of SE gene induced by 6 brands of RJ (2: RJ, Fengzhiyu bio, Hangzhou, China; 3: RJ, Yiyuan bio, Beijing China; 4: RJ, Baosheng bio, Guangzhou, China; 5: RJ, Wang’ s bio, Jiangxi, China; 6: RJ, Laoshan bio, Nanjing, China; 7: RJ, Leishi bio, Shanghai, China) which collected from different region. RT-PCR analysis shows that the expression of SE which induced by 2 and 5 have significantly difference (p < 0.05).

The quality of RJ merchandises varies due to the place of origin, bee species and pollen collection. Therefore, we tested several brands of RJ at 5 mg/mL. As shown in Fig 1b, SE expression was affected by the origin of RJ suggesting that these RJ products may contain different ingredients. All RT-PCR experiments had been repeated six times.

### Analysis of active ingredients of RJ

To determine the responsible active ingredient of RJ for SE down-regulation, we chose the brand produced in Hangzhou China for fractionation. Firstly, we used semi permeable membrane dialysis to separate RJ solution into two categories: above and below 10kDa. As shown in Fig 2A, the fraction < 10kDa had a similar effect as the RJ before the purification in SE reduction. We further separated the portion of < 10kDa molecular weight into fat-soluble components and water-soluble components using methanol. The result shows that the water-soluble components of < 10 kDa had the same effect as the original RJ in SE reduction (Figure 2B).

**Fig. 2.**
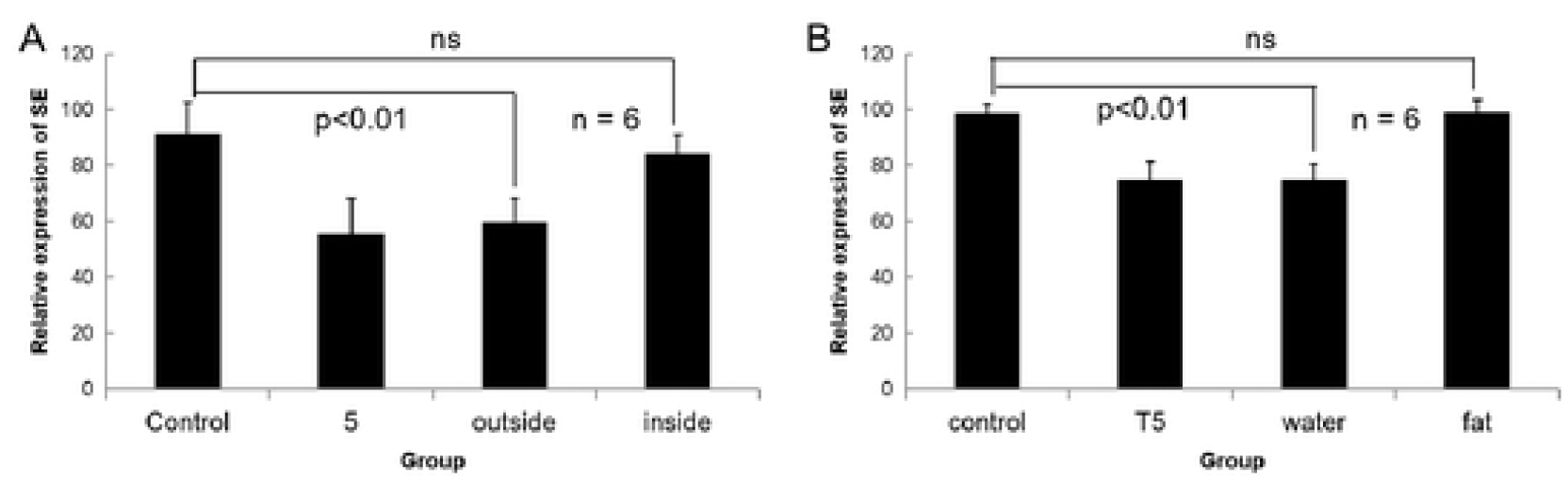
Separated RJ component effect the gene expression of SE in HepG2 cells. (A) Representative Real-time polymerase chain reaction (RT-PCR) showing the expression changes of SE in the presence of 5 (RJ concentration: 5 mg/mL), its outside solution and inside solution of semipermeable membrane. Outside solution was the fraction which molecular weight less than 10kDa; inside solution was the portion which larger than 10 kDa. Vertical axis represent as the value of SE expression divided by the value of GAPDH expression. Horizontal axis represent as categories of sample. Control group was added growth medium only. RT-PCR analysis shows that 5 and outside solution of semipermeable membrane significantly decrease mRNA expression of SE in HepG2 cells (p < 0.01) but not inside solution. (B) The results of RT-PCR showing the expression of SE gene affected by 5 (RJ concentration: 5 mg/mL), water-soluble component (water) and fat-soluble component (fat) of outside solution of semipermeable membrane (separated by ethanol). RT-PCR analysis shows that the expression of SE which induced by water-soluble component have significantly difference (p < 0.05) as same as RJ.

## Discussion

As the Chinese economy has experienced an accelerating growth of economy in recent years, the prevalence of obesity and the incidence of fatty liver disease has gone up along with it. Therefore, early detection and treatment are the only way to the prognosis of the disease. Although steady progress has been made in elucidating the pathogenesis, still there is no specific therapeutic drugs for the disease currently. Identifying therapeutic targets and advancing drug development are urgent. For example, NAFLD is also known as metabolic-related fatty liver disease (MAFLD), which is related to metabolic syndrome. Previous studies indicated that excessive accumulation of cholesterol plays an important role in the fatty liver disease ^[14]^. Taking control of lipid metabolism can be an important way to prevent and cure the fatty live disease. Today, STATINS are the most commonly used drugs, which can reduce cholesterol production in liver for prevention purposes and treatment of fatty live disease. But some studies have shown that STATINS do not have a clear effect on preventing and reversing liver fibrosis and it has serious side effects such as elevating liver enzymes, gastrointestinal reactions, and rhabdomyolysis. To explore new treatment methods and discover a new drug for fatty liver disease is necessary. Royal Jelly as a kind of natural food can be eaten directly and absorbed. There are sufficient supplies in the market and it is convenient to purchase. Previous studies have demonstrated a reduction in SE expression in RJ fed mice ^[23]^. However, a mouse is a complex biological system. Conclusions that are solely derived from animal study need to be elucidated at the molecular and cellular levels. That is where we came in. We used cell experiments to further clarify the role of RJ and explore the active ingredients in RJ. We added RJ to HepG2 cells, the expression of SE gene decreased significantly, which is comparable to the previous animal experiment. Interestingly, as the concentration of RJ increased, the decrease of SE gene expression was kept around 50%, indicating that RJ only reduces SE expression to certain degree and does not terminate the synthesis of cholesterol. This is good because the cholesterol is an essential component of the cell membrane and without cholesterol a cell cannot survive. RJ can prevent the excessive production of cholesterol in the liver, and therefore, it can be a new way to prevent and treat the fatty liver disease.

The color, taste and the chemical content of RJ can be affected by the species diversity of bees as well as the plants where bees live. In our study, we tested the RJ from seven different regions of China and found the SE reduction among them were differ. Fractionation after fractionation, we found the active ingredient is in the water-soluble fraction of the small molecules (molecular weight < 10kDa). We speculate that it is likely to be trans-10-hydroxy-2-decenoic acid (10-HDA), which is an unsaturated fat acid with low molecular weight, fitting the characteristics. 10-HDA also a marker to evaluate the quality of RJ, because 10-HDA is found only in Royal Jelly.

Taken together, Royal Jelly can directly act on the liver cells by decreasing rather than terminating cholesterol production in the liver. Therefore, RJ has great potential to be a new way to prevent and treat fatty liver disease induced by cholesterol accumulation. Its active component is a small water-soluble molecule that down-regulates SE expression. If this component is 10-HDA, we can extract this substance from RJ for future experiments, and further explore the specific pathway of the SE inhibition.

## Conclusion

RJ down-regulates SE expression in HepG2 cells (Fig 3). Its active ingredient for this function is a small water-soluble component (< 10kDa) likely to be 10-HDA.

**Fig. 3.**
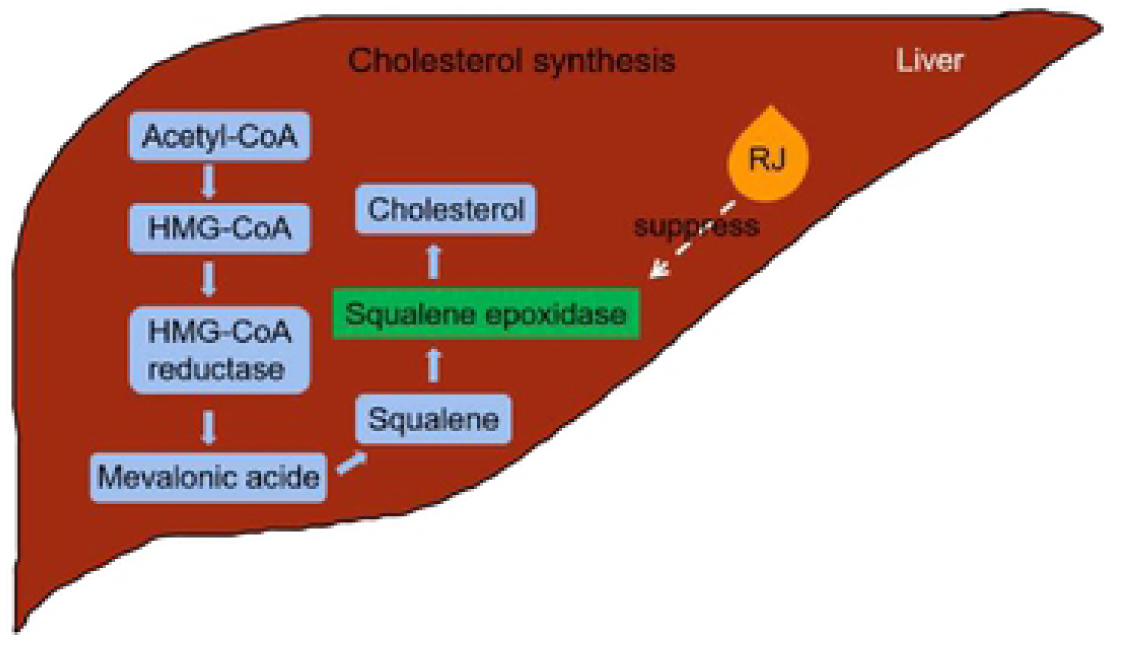
Mechanism of suppression gene expression of cholesterol by RJ. RJ decreases gene expression of squalene epoxidase (SE), a key enzyme in cholesterol biosynthesis. Thus, RJ might lower cholesterol in liver by decreasing SE.

